# RDC complex executes a dynamic piRNA program during *Drosophila* spermatogenesis to safeguard male fertility

**DOI:** 10.1101/2020.08.25.266643

**Authors:** Peiwei Chen, Yicheng Luo, Alexei A. Aravin

## Abstract

piRNAs are small non-coding RNAs that guide the silencing of transposons and other targets in animal gonads. In *Drosophila* female germline, many piRNA source loci dubbed ‘piRNA clusters’ lack hallmarks of active genes and exploit an alternative path for transcription, which relies on the Rhino-Deadlock-Cutoff (RDC) complex. It remains to date unknown how piRNA cluster transcription is regulated in the male germline. We found that components of RDC complex are expressed in male germ cells during early spermatogenesis, from germline stem cells (GSCs) to early spermatocytes. RDC is essential for expression of dual-strand piRNA clusters and transposon silencing in testis; however, it is dispensable for expression of Y-linked *Suppressor of Stellate* piRNAs and therefore *Stellate* silencing. Despite intact *Stellate* repression, *rhi* mutant males exhibited compromised fertility accompanied by germline DNA damage and GSC loss. Thus, piRNA-guided repression is essential for normal spermatogenesis beyond *Stellate* silencing. While RDC associates with multiple piRNA clusters in GSCs and early spermatogonia, its localization changes in later stages as RDC concentrates on a single X-linked locus, *AT-chX*. Dynamic RDC localization is paralleled by changes in piRNA cluster expression, indicating that RDC executes a fluid piRNA program during different stages of spermatogenesis.

## INTRODUCTION

Transposable elements (TEs) are selfish genetic elements that have the ability to propagate in the genome. When unchecked, transposition of TEs can cause overwhelming DNA damage and, eventually, genome instability. This poses a particular threat to germ cells, and TE de-repression often leads to reproductive defects like sterility. To cope with this, a small RNA-mediated genome defense mechanism involving PIWI proteins and PIWI-interacting RNAs (piRNAs) is employed in animal gonads to silence TEs (Ozata et al., 2019).

In ovaries of *Drosophila melanogaster*, most piRNAs are made from so-called dual-strand piRNA clusters, where both genomic strands are transcribed to give rise to piRNA precursors. Transcription of dual-strand piRNA clusters is unusual in a number of ways: 1) there is no clear promoter signature for transcription initiation (Mohn et al., 2014), 2) splicing, termination and poly-adenylation of nascent transcripts are all suppressed (Chen et al., 2016; Mohn et al., 2014; Zhang et al., 2014), 3) transcription occurs at the presence of H3K9me3 (Rangan et al., 2011), a histone modification generally seen as a repressive mark for gene expression. Prior work has shown that non-canonical transcription and co-transcriptional processing of piRNA precursors depend on the RDC complex, composed of Rhino (Rhi), Deadlock (Del) and Cutoff (Cuff) proteins, that bind dual-strand clusters (Mohn et al., 2014; Zhang et al., 2014). Rhi belongs to the HP1 family and binds H3K9me3 through its chromo-domain, anchoring the RDC complex onto dual-strand clusters (Le Thomas et al., 2014; Mohn et al., 2014; Zhang et al., 2014). Cuff is a homolog of the conserved cap-binding protein Rai1 that was reported to suppresses both splicing (Zhang et al., 2014) and transcriptional termination (Chen et al., 2016), in order to facilitate the production of long, unspliced piRNA precursors. Del, on the other hand, recruits a paralog of transcription initiation factor TFIIA-L, Moonshiner (Moon), to initiate transcription in hostile heterochromatin environment (Andersen et al., 2017). Together, the RDC complex conveys transcriptional competence to dual-strand piRNA clusters, and the majority of piRNA production collapses when one of its components is missing.

While piRNA pathway is known to be active in both male and female germline of *Drosophila*, expression and functions of Rhi, Del and Cuff were studied exclusively during oogenesis. Mutations of *rhi*, *del* and *cuff* were shown to cause female sterility, however, mutant males remained fertile (Berg and Spradling, 1991; Schüpbach and Wieschaus, 1991). In addition, *rhi*, *del* or *cuff* are predominantly expressed in ovaries, with low or no expression in testes and somatic tissues (Brown et al., 2014; Vermaak et al., 2005; Volpe et al., 2001). Similarly, Moon was also believed to be ovary-specific, given that a high expression level could only be found in ovaries (Andersen et al., 2017). Finally, the silencing of *Stellate* by abundant *Suppressor of Stellate* piRNAs was shown to be unperturbed in testes of *rhi* mutants, suggesting that *rhi* is not required for piRNA biogenesis in males (Klattenhoff et al., 2009). Collectively, these findings led to the notion that RDC complex is dispensable for piRNA pathway in male germline, raising the question of how piRNA cluster expression is controlled during spermatogenesis.

Here, we describe a developmentally regulated assembly of RDC complex in testes. We found that low expression of RDC complex components can be attributed to the fact that only a small subset of cells at early stages of spermatogenesis express *rhi*, *del* and *cuff*. Loss of RDC complex in testes results in a collapse of piRNA production, global TE de-repression, and, ultimately, compromised male fertility, supporting an indispensable role of RDC complex in spermatogenesis. Even though RDC complex is assembled and functional in both sexes, we found differential genome occupancies of RDC complex between two sexes, correlating with sexually dimorphic usage of genome-wide piRNA source loci. Finally, RDC complex appears to exhibit dynamic binding on different piRNA clusters during spermatogenesis, allowing different piRNAs source loci to be used at different stages of early sperm development.

## RESULTS

### Rhi is required for normal male fertility

Previous studies showed that, while *rhi* is required for female fertility, it is dispensable for male fertility (Klattenhoff et al., 2009; Volpe et al., 2001). In agreement with this, we found that *rhi* mutant males indeed produce progenies when crossed with wildtype females. However, careful examination of the male fertility by sperm exhaustion test (Sun et al., 2004) revealed significantly compromised fertility in *rhi* mutant males. Even though most *rhi* mutant males were initially fertile, the percentage of fertile males dropped as they aged and stayed low from day 3 in comparison to heterozygous sibling controls (Figure 1A). To probe male fertility more quantitatively, we repeated the test and counted numbers of progenies for each male every day. We found that even young, 1-day-old *rhi* mutant males, which were fertile, produced fewer progenies than heterozygous sibling controls (Figure 1A). Also, *rhi* mutant males produced nearly no progeny after two-day sperm exhaustion, while heterozygous sibling controls continued to produce ~100 progenies on average throughout the sperm exhaustion process. These results demonstrate that male fertility is substantially compromised at the absence of *rhi*, suggesting an indispensable role of *rhi* in maintaining normal male fertility.

**Figure 1.**
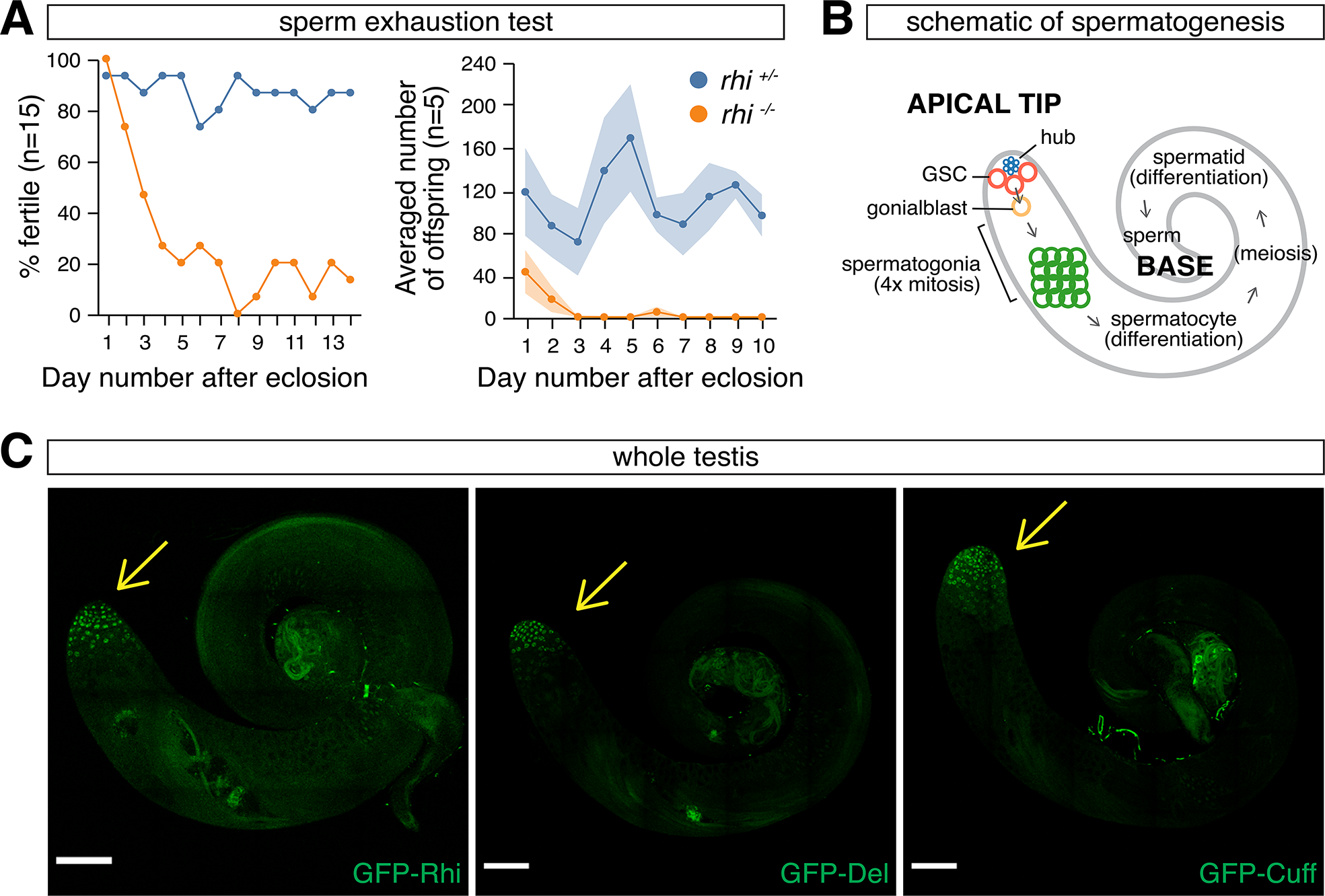
Rhi is required for normal male fertility, and components of RDC complex are expressed at the apical tip of testis. (A) Compromised fertility *of rhi* mutant males. Sperm exhaustion test of *rhi*^*2/KG*^ mutant (orange) and heterozygous sibling control (blue) males. Left: percentages of fertile males 1-14 days post-eclosion (n=15). Right: averaged numbers of offspring per male 1-10 days after eclosion (n=5). Shaded areas display standard error. Two charts report results from two independent sperm exhaustion tests. (B) A schematic of spermatogenesis, showing major developmental stages of male germline as well as the somatic hub that serves as germline stem cell (GSC) niche. (C) Expression of GFP-tagged Rhi (left), Del (middle) and Cuff (right), driven by their respective promoters. Expression of all three proteins can only be seen at the apical tip of testis (pointed to by the yellow arrow). Scale bar: 100µm.

### RDC complex is assembled in nuclei of germ cells from GSCs to early spermatocytes

The dependency of normal male fertility on *rhi* prompted us to re-examine whether Rhi-Del-Cuff (RDC) complex is assembled in testis. modENCODE data and previous work showed that tissue-wide mRNA levels of *rhi, del* and *cuff* are high in ovaries but low in testes and the soma (Brown et al., 2014; Vermaak et al., 2005; Volpe et al., 2001), which led to the notion that RDC might be ovary-specific (Vermaak et al., 2005; Vermaak and Malik, 2009; Volpe et al., 2001). To examine expression of Rhi, Del and Cuff in testis, we took an imaging-based approach that provides single-cell resolution and preserves spatial information. We examined expression of individual components of RDC complex using GFP-tagged Rhi, Del and Cuff driven by their native promoters. All three proteins are expressed at the apical tip of testis that contain germ cells at early steps of spermatogenesis (Figure 1B and 1C), indicating that all three components of RDC complex are expressed in testis, though only in a subset of the cells.

Rhi, Del and Cuff form foci in nuclei (Figure 2A). To test whether all three proteins co-localize in nuclear foci, we tagged Rhi by a different fluorophore and analyzed its localization with Del and Cuff. Indeed, Rhi co-localizes with both Del and Cuff (Figure 2A), consistent with the formation of RDC complex. Next, we examined inter-dependence of Rhi, Del and Cuff localization (Figure 2B). In either *del* or *cuff* mutants, Rhi becomes dispersed and no longer forms puncta in nuclei. Similarly, Del also disperses at the absence of Rhi or Cuff. Expression of Cuff is strongly decreased in both *rhi* and *del* mutants, indicating its destabilization. Therefore, Rhi, Del and Cuff co-localize in distinct nuclear foci that depend on the simultaneous presence of all three proteins.

**Figure 2.**
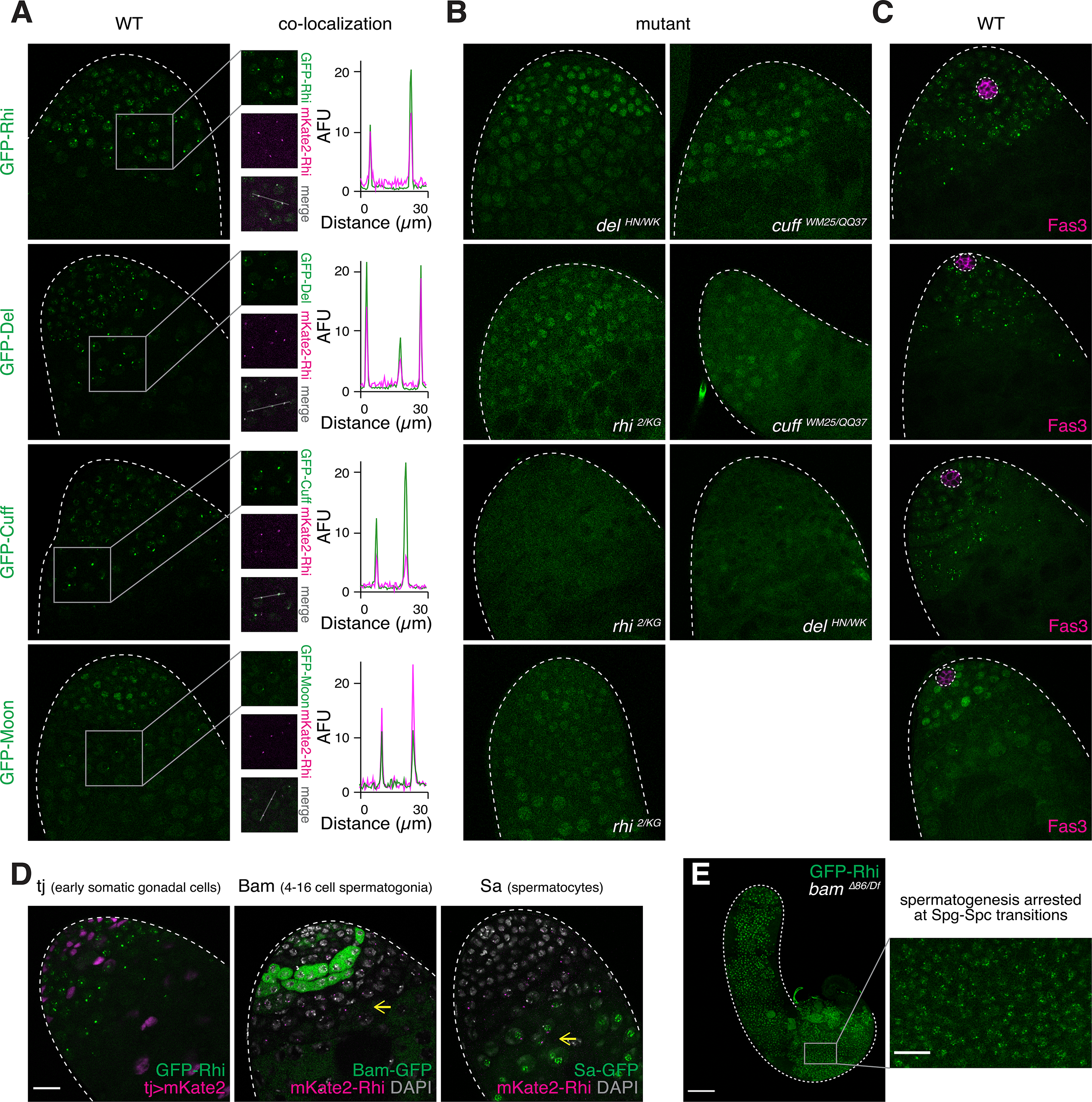
RDC complex is assembled in early male germ cells. (A) Rhi, Del, Cuff and Moon co-localize in nuclear foci. Confocal images showing apical tips of testes expressing GFP-tagged Rhi, Del, Cuff and Moon (top to bottom), driven by their native promoters. Co-localization with mKate2-tagged Rhi in nuclear foci are shown on the right. Signal intensities along the marked line are plotted for each of four co-localization analysis. AFU, arbitrary fluorescence units. (B) Inter-dependence of Rhi, Cuff, and Del localization in nuclear foci, as well as the dependence of Moon localization on Rhi. Confocal images showing apical tips of testes expressing GFP-Rhi in *de*l^*HN/WK*^ and *cuff*^*WM25/QQ37*^, GFP-Del in *rhi*^*2/KG*^ and *cuff*^*WM25/QQ37*^, GFP-Cuff in *rhi*^*2/KG*^ and *del*^*HN/WK*^, and GFP-Moon in *rhi*^*2/KG*^ mutant backgrounds. Nuclear foci of each protein dispersed or disappeared in respective mutants. (C) Rhi, Del, Cuff and Moon are expressed in GSCs. Immuno-fluorescence of testes expressing GFP-tagged Rhi, Del, Cuff and Moon, driven by their respective promoters, stained for somatic hub marker Fas3. Note that GSCs directly adjacent to Fas3-positive hub express all four proteins. (D) Rhi is not expressed in somatic gonadal cells, and its germline expression continues beyond spermatogonia till early spermatocytes. Confocal images showing fluorescently tagged Rhi driven by its own promoter with somatic gonadal cells marked by *tj-Gal4>UASp-mKate2* (left), 4-16 cell spermatogonia marked by Bam-GFP (middle) and spermatocytes marked by Sa-GFP (right). Expression of Rhi in early spermatocytes is pointed to by yellow arrows. (E) Rhi expression upon arrest of spermatogenesis in *bam* mutants. Confocal image showing *bam*^∆*86/Df*^ mutant testis expressing GFP-Rhi driven by *rhi* promoter, where spermatogenesis is arrested at the spermatogonia-to-spermatocyte transition stage. Note that spermatogonia are expanded and virtually all germ cells express Rhi. An enlarged view of the basal part of mutant testis is shown at the bottom, which shares scale bar with (D). All images share scale bars with (D), except for (E). Scale bars: 20µm (D) and 100µm (E).

We further characterized the expression of RDC complex in testis. Rhi, Del and Cuff are expressed in nuclei of germline stem cells (GSCs) that are directly adjacent to somatic hub cells labeled by Fas3, but not in hub cells (Figure 2C). Rhi expression continues beyond spermatogonia marked by Bam, until early spermatocytes that express Sa (Figure 2D). In *bam* mutant testes, where spermatogenesis is arrested at the spermatogonia-to-spermatocyte transition stage, we observed an expansion of spermatogonia and expression of Rhi throughout entire testes (Figure 2E). In addition to germ cells and hub cells, the apical tip of testes contains somatic gonadal cells (cyst stem cells and early cyst cells) that can be distinguished from germ cells by Tj expression. Rhi is not expressed in somatic cells that express Tj (Figure 2D), confirming its restriction to the germline. Taken together, we conclude that RDC complex is assembled in male germ cells during spermatogenesis, from GSCs to spermatogonia and early spermatocytes.

In ovaries, RDC complex is known to promote piRNA cluster transcription by two mechanisms: 1) suppression of premature transcriptional termination, a function mediated by Cuff (Chen et al., 2016), and 2) licensing of non-canonical transcriptional initiation, a function that requires the recruitment of a basal transcriptional factor TFIIA-L paralog, Moonshiner (Moon) (Andersen et al., 2017). Expression of Moon was reported to be specific to female germline, raising the question of whether RDC complex can fulfill its function in male germline, if its functional partner is missing. However, we observed Moon expression in testis using GFP-tagged Moon driven by its native promoter (Figure 2A). Importantly, Moon is expressed at similar stages as components of RDC complex, from GSCs to spermatogonia and early spermatocytes at the apical tip of testis (Figure 2A and 2C). Furthermore, Moon co-localizes with Rhi in nuclear foci, and its focal localization is abolished in *rhi* mutants (Figure 2A and 2B). On the contrary, Rhi localization appears normal in two different *moon* mutants, *moon*^∆*1*^ and *moon*^∆*28*^, indicating that it acts genetically downstream of RDC complex (Figure S1). These observations suggest that RDC complex can recruit Moon to license transcription initiation in the male germline.

### Loss of RDC complex causes DNA damage and germ cell death in testis

To identify cellular mechanisms underlying fertility decline in males lacking RDC complex, we examined morphology of germ cells marked by Vasa-GFP in *rhi* mutants. Normally, Vasa-positive germ cells are tightly packed at the apical tip of testis, as we observed in testes of heterozygous control males (n=162; Figure 3A). However, half of *rhi* mutant testes (54.9%, n=134/244) had visibly fewer germ cells with prominent gaps in between, indicative of an elevation of germ cell death (Figure 3A). Furthermore, another quarter of *rhi* mutant testes (25.4%, n=62/244) completely lost germ cells altogether, and only 19.7% (n=48/244) showed wildtype-like germline morphology (Figure 3A and 3B). We concluded that loss of RDC complex leads to a reduction in the germ cell count in testis.

**Figure 3.**
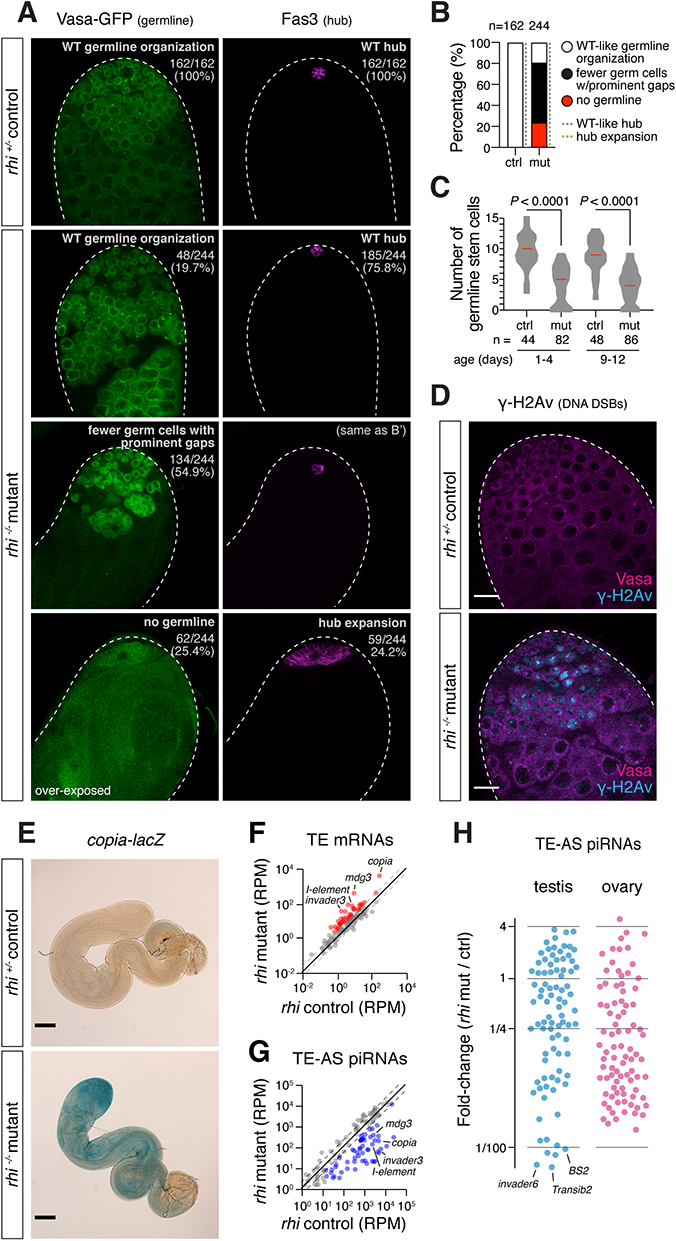
Loss of Rhi causes germ cell death, DNA damage and TE de-repression. (A) Loss of germ cells in testes of *rhi* mutants. Left: expression of germ cell marker, Vasa-GFP driven by *vasa* promoter, in testes of *rhi*^*2/KG*^ mutants and heterozygous control. Right: somatic hub cells that form a niche for GSCs are marked with Fas3. Classification of germ cell phenotype (defined by number and organization of germ cells that express Vasa) and hub size phenotype (defined by cells that express Fas3) are labeled at the top right with corresponding statistics. All images on the same scale as (D). (B) Quantification of germ cell survival and hub size in *rhi* heterozygous control (left) and mutant (right) testes shown in (A). n, number of testes examined. (C) Loss of GSCs in testes of *rhi* mutants. Violin plot showing GSC numbers in *rhi*^*2/KG*^ mutant and age-matched heterozygous sibling control testes. Median of GSC number is marked red. GSC number is acquired by counting the total number of Vasa-positive cells directly adjacent to Fas3-positive hub in 3D. *P* < 0.0001 based on Mann–Whitney–Wilcoxon test. n, number of testes counted for each genotype and age group. (D) Accumulation of DNA double-strand breaks (DSBs) in male germline of *rhi*^*2/KG*^ mutants. Immuno-fluorescence of *rhi* heterozygous control and mutant testes, stained for γ-H2Av, a marker for DNA DSBs, and Vasa, a germline marker. Scale bars: 20µm. (E) De-repression of *copia* reporter in testes of *rhi* mutants. Brightfield images showing *rhi* heterozygous control and *rhi*^*2/KG*^ mutant testes expressing *copia-lacZ*, after X-gal staining. *copia* LTR containing its promoter is fused upstream to *lacZ* gene. Scale bar: 100µm. (F) De-repression of TEs in testes of *rhi* mutants measured by polyA+ RNA-seq. Scatter plot showing expression of TE mRNAs in *rhi*^*2/KG*^ mutant versus heterozygous control testes. TEs that show ≥2-fold de-repression and ≥1 RPM averaged levels are marked red. (G) Loss of TE-targeting piRNAs in testes of *rhi* mutants. Scatter plot showing expression of TE-antisense piRNAs in *rhi*^*2/KG*^ mutant versus heterozygous control testes. TE-antisense piRNAs that show ≥2-fold reduction and ≥10 RPM averaged levels are marked blue. (H) Loss of TE-targeting piRNAs in *rhi* males and females. Scatter plot showing fold-change of TE-antisense piRNAs upon loss of *rhi* in testis (left, blue) and ovary (right, pink). Note that piRNAs targeting several TE families demonstrate over 100-fold reduction in *rhi* testis, a magnitude not observed in ovary.

Next, we examined impacts of Rhi loss on the resident germline stem cell (GSC) population. We quantified the number of GSCs per testis by counting the number of Vasa-positive germ cells directly adjacent in 3D to the somatic GSC niche labeled by Fas3. We found a reduction of GSCs in testes of *rhi* mutants compared with heterozygous controls in two age groups (1-4 and 9-12 days old) (*P*<0.0001, Mann–Whitney–Wilcoxon test, Figure 3C). About a quarter of *rhi* mutant testes did not have any GSC at all. Accordingly, we observed an expansion of Fas3-positive hub at a similar rate (24.4%, n=59/244), which usually occurs at the absence of GSCs and is never seen in control testes (Figure 3A). Hence, GSC population sizes shrink drastically in testes lacking RDC complex. Staining testes for the phosphorylated H2A variant (γ-H2Av), a marker for DNA double-strand breaks (DSBs), revealed massive accumulation of unrepaired DNA DSBs in early germ cells of *rhi* mutant testes (Figure 3D). Unrepaired DNA DSBs likely causes germ cell death in *rhi* mutants. Overall, our results suggest that the loss of germ cells, including GSCs, accompanied by widespread unrepaired DNA DSBs is responsible for the compromised fertility of *rhi* mutant males.

### RDC complex is required for TE silencing in testis

Widespread DNA DSBs can result from TE transposition. To quantify TE expression, we sequenced polyadenylated (polyA+) RNAs from *rhi* mutant and control testes. PolyA+ RNA-seq revealed 36 TE families showing more than 2-fold up-regulation in testes of *rhi* mutants (Figure 3F). Among them, the most de-repressed ones include *mdg3* (33-fold), *copia* (13-fold) and *invader 3* (12-fold). To verify TE de-repression, we employed a *copia*-lacZ reporter, where the long terminal repeat (LTR) of *copia* containing *copia* promoter is fused upstream to the *lacZ* gene and its expression can be directly examined by X-gal staining (Kalmykova et al., 2005). *copia* is known to be active in the male germline (Pasyukova et al., 1997) and has the highest expression level among all TEs in testes (see accompanying manuscript). Whereas no detectable X-gal staining was seen in control testes, robust staining was observed in *rhi* mutants (Figure 3E), confirming strong de-repression of *copia* at the absence of *rhi*. These results show that TEs are de-repressed in testes lacking a functional RDC complex.

In ovaries, RDC complex is required for piRNA production from dual-strand clusters to ensure efficient TE silencing (Andersen et al., 2017; Chen et al., 2016; Mohn et al., 2014; Zhang et al., 2014). To examine piRNA biogenesis, we sequenced and analyzed small RNAs in *rhi* mutant and control testes. We observed a loss of antisense piRNAs targeting many TE families in *rhi* mutant testes (Figure 3G), suggesting an overall defect in piRNA production. For TEs showing strong up-regulation in *rhi* mutant testes, we observed a concurrent loss of antisense piRNAs (e.g., *mdg3, copia* and *invader3*). Notably, there are antisense piRNAs against several TE families (e.g., *BS2*, *Transib2*, *invader6*) that show over 100-fold reduction in *rhi* mutant testes, a magnitude not observed for any TE family in *rhi* mutant ovaries (Figure 3H). These results show that efficient production of TE-silencing piRNAs in testis depends on the RDC complex, without which many TEs are de-repressed, causing DNA damage and germ cell death in testis.

### RDC complex is required for piRNA production from dual-strand clusters in early male germ cells

To understand the role of RDC complex in piRNA cluster expression in testis, we analyzed effects of *rhi* mutation on piRNA production from major piRNA clusters. The loci that generate piRNAs in testis were *de novo* defined leading to identification of several novel piRNA clusters (see accompanying manuscript). Rhi was dispensable for expression of uni-strand piRNA clusters, *flam* and *20A* (Figure 4A), similar to results from ovary studies (Klattenhoff et al., 2009; Mohn et al., 2014; Zhang et al., 2014). Surprisingly, we found that Rhi was also dispensable for piRNA production from the Y-linked *Su(Ste)* locus, which is the most active piRNA cluster in testis (see accompanying manuscript). Unlike *flam* and *20A*, *Su(Ste)* is a dual-strand cluster that generates piRNAs from both genomic strands. We confirmed by RNA fluorescence *in situ* hybridization (FISH) that piRNA precursor transcription from *Su(Ste)* appeared intact in testes without Rhi (Figure 4C and 4D). In contrast to *Su(Ste)*, piRNA production from other major dual-strand clusters, including the Y-linked *h17* cluster, collapses in *rhi* mutant testes (Figure 4A), indicating that expression of the majority of dual-strand piRNA clusters in testis relies on Rhi. Interestingly, dependence of dual-strand cluster expression on Rhi varies between sexes: *38C* is more affected by loss of *rhi* than *42AB* in testis, while the opposite is found in ovary. Furthermore, piRNA production from both strands of complex satellites, which we recently found to behave as dual-strand piRNA clusters (see accompanying manuscript), also drastically declined in *rhi* mutant testes and ovaries (Figure 4B). These results show that RDC complex is essential for piRNA production from a large fraction of piRNA clusters in male germline.

**Figure 4.**
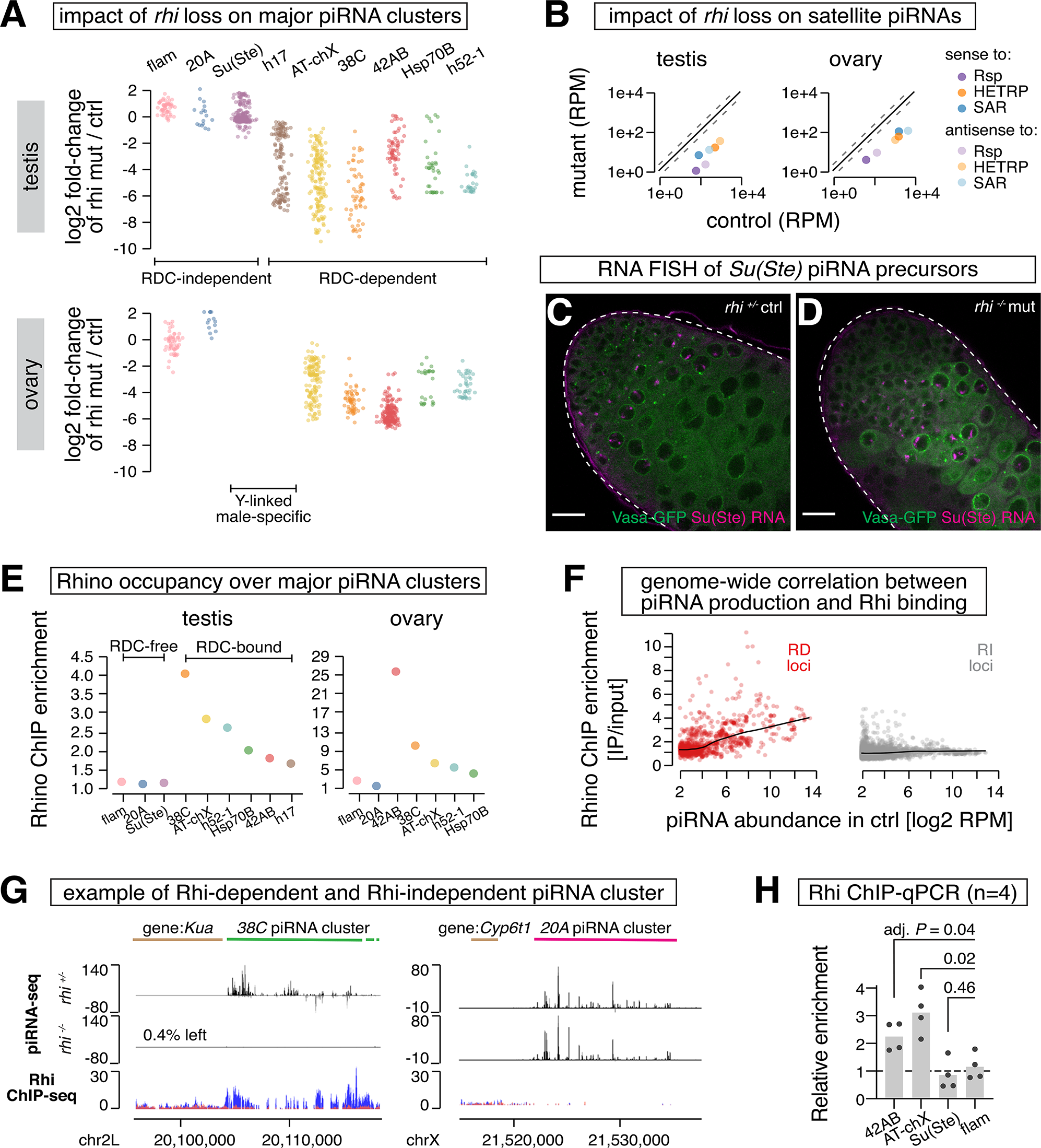
RDC complex is required for piRNA production from dual-strand piRNA clusters. (A) Impacts of *rhi* loss on piRNA production from major piRNA clusters in testis and ovary. Scatter plot showing fold-change of piRNA production from 1Kb genomic windows spanning major piRNA clusters in testis (top) and ovary (bottom), upon loss of *rhi*. Two Y-linked, male-specific clusters are not present in female genome. Each cluster is given a unique color. (B) Impact of *rhi* loss on piRNA production from satellite repeats in testis and ovary. Scatter plot showing levels of complex satellite-mapping piRNAs in *rhi*^*2/KG*^ mutant and control testis (left) and ovary (right). Each complex satellite is assigned a color, and piRNAs sense to satellite consensus are marked with higher opacity than antisense ones. (C-D) Unperturbed expression of *Su(Ste)* piRNA precursors in testes of *rhi* mutants. RNA fluorescence *in situ* hybridization of *Su(Ste)* piRNA precursors in *rhi* control (C) and *rhi*^*2/KG*^ mutant (D) testes that express Vasa-GFP. Scale bar: 20µm. (E) Rhi occupancy over major piRNA clusters in testes and ovary. Scatter plot showing Rhi ChIP-seq enrichment over major piRNA clusters in testis (left) and ovary (right). Shown are averages of two biological replicates. Two Y-linked, male-specific clusters are not present in female genome. Each cluster is colored the same way as in (A). (F) Scatter plot showing the relationship between Rhi ChIP enrichment and piRNA production over 1Kb genomic windows. Loci that show ≥4-fold decline in piRNA production at the absence of *rhi* are defined as Rhi-dependent (“RD”, red), otherwise Rhi-independent (“RI”, gray). Black lines show local regression. (G) Examples of Rhi-dependent (*38C*, left) and Rhi-independent (*20A*, right) piRNA clusters. piRNA-seq in *rhi* mutant and control testes are shown at the top, and Rhi ChIP-seq is shown at the bottom (blue: IP, red: input). (H) Rhi does not bind *Su(Ste)* cluster in testis. Bar graphs showing Rhi ChIP-qPCR (n=4) over four piRNA clusters in testis. Adjusted *P-*values are from multiple t-tests corrected for multiple comparisons by the Holm-Sidak method. Uni-strand piRNA cluster *flam* not bound by Rhi serves as a negative control.

Since RDC complex forms distinct foci in the nuclei of germ cells (Figure 2A), we set out to test if Rhi binds the chromatin of dual-strand clusters whose expression depends on RDC complex, as reported in ovary (Klattenhoff et al., 2009; Mohn et al., 2014; Zhang et al., 2014). Given that expression of RDC complex is restricted to a small number of cells at the apical tip of testis, we used *bam* mutant testes, where Rhi-expressing spermatogonia are expanded (Figure 2E), to perform ChIP-seq of Rhi. All major dual-strand clusters, with an exception of *Su(Ste)*, were enriched for Rhi binding (Figure 4E). In agreement with ChIP-seq, independent ChIP-qPCR showed no evidence of Rhi binding on *Su(Ste)* locus (n=4) (Figure 4H). Rhi was also absent on chromatin of uni-strand clusters, *flam* and *20A*. Importantly, the binding of Rhi on different loci correlates with its effect on promoting piRNA cluster expression. Rhi does not bind, and is dispensable for piRNA production from, uni-strand clusters and *Su(Ste)*, while it binds, and is required for expression of, other dual-strand clusters (Figure 4A and 4E). Also, dual-strand clusters that show the highest levels of overall Rhi binding, *38C* and *AT-chX*, demonstrate the strongest Rhi dependence for piRNA production. To characterize the relationship between Rhi binding and piRNA production on a genome-wide scale, we analyzed Rhi binding and piRNA production in 1Kb genomic windows spanning the entire genome (Figure 4F). For loci that depend on Rhi to produce piRNAs, we observed a correlation between Rhi binding and piRNA levels. On the other hand, loci that continue to produce piRNAs at the absence of Rhi usually have little, if any, Rhi binding. Collectively, our data indicate that RDC complex physically binds the chromatin and ensures the expression of dual-strand piRNA clusters, with a notable exception of *Su(Ste)*.

### Sexually dimorphic genome occupancy of RDC complex sculpts sex-specific piRNA program

piRNA profiles are distinct in male and female gonads, and expression of dual-strand piRNA clusters are sexually dimorphic (see accompanying manuscript). To explore if RDC complex might be involved in orchestrating sex-specific piRNA programs, we profiled Rhi binding on the genome in ovaries under identical ChIP-seq conditions as in testes. This analysis revealed differences in Rhi genome occupancy between sexes among top piRNA clusters (Figure 5A). For example, Rhi is more enriched on *38C* than *42AB* in testes, whereas the reciprocal is seen in ovaries, correlating with differential piRNA production from these two loci in two sexes. In addition, *80EF* and *40F7* clusters have high levels of Rhi binding in ovary but low in testis, mirrored by abundant piRNA production from these two loci in ovary but not in testis. Finally, an ovary-specific dual-strand piRNA cluster, *Sox102F*, is bound by Rhi in ovary, while there is no evidence of Rhi binding at *Sox102F* in testis where it is inactive (Figure 5B). Altogether, the observed link between Rhi binding and piRNA production between males and females suggests that the sex-specific Rhi binding on piRNA clusters is responsible for sculpting a sexually dimorphic piRNA program.

**Figure 5.**
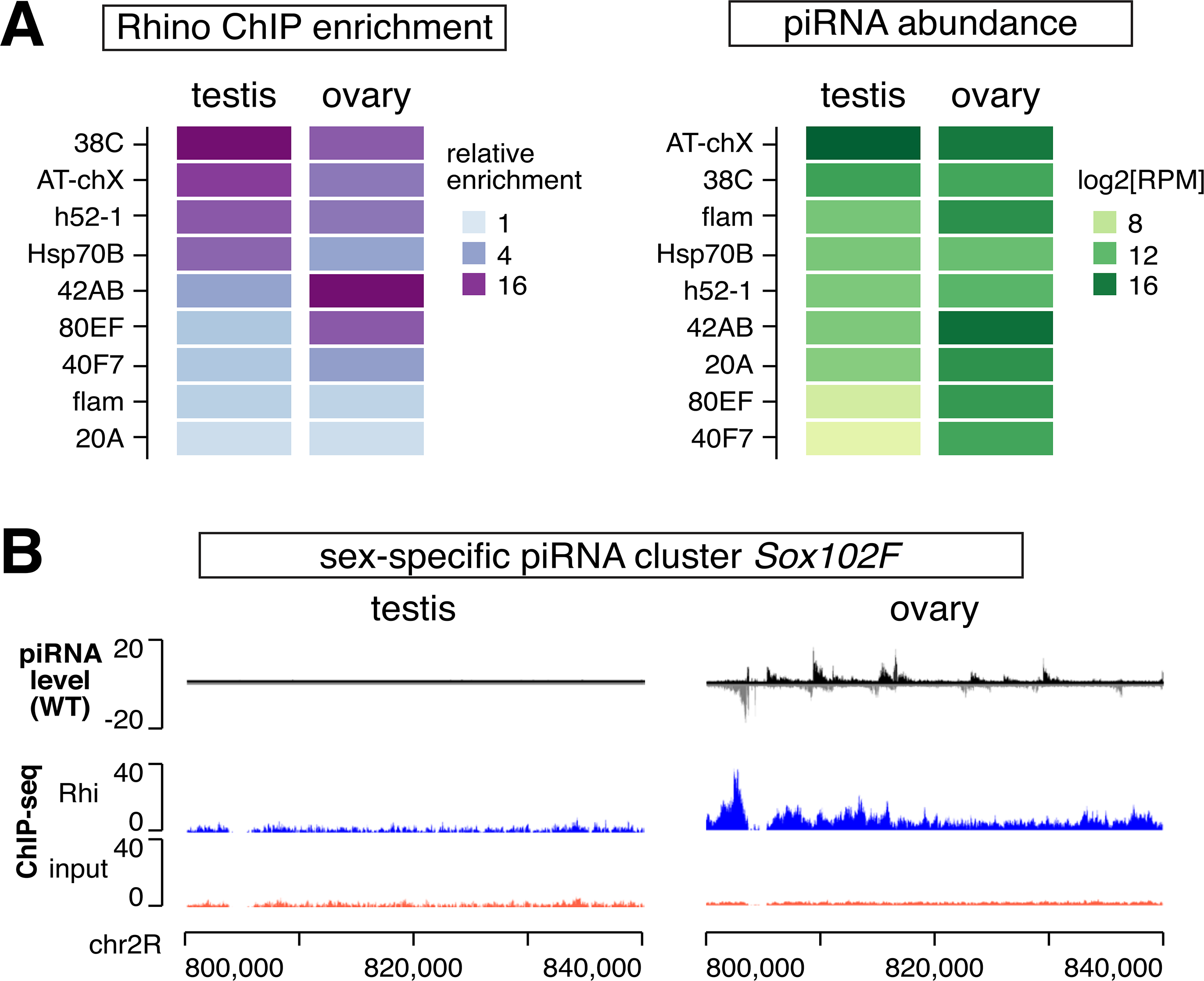
Sexually dimorphic RDC genome occupancy sculpts sex-specific piRNA program. (A) Heatmaps showing relative enrichment of Rhi binding over major piRNA clusters in two sexes determined by ChIP-seq. (B) Heatmaps showing piRNA production from major piRNA clusters in two sexes. (C) *Sox102F* piRNA cluster produces piRNAs exclusively in ovary. Rhi is enriched on *Sox102F* in ovary, but not in testis.

### RDC complex enables dynamic piRNA production during spermatogenesis

ChIP-seq provides the genome-wide profile of Rhi binding, but it masks possible differences of Rhi localization among individual cells. Imaging of Rhi revealed distinct Rhi localization in nuclei of germ cells at different stages of spermatogenesis (Figure 6A). In the nuclei of GSCs and spermatogonia, Rhi forms many discrete foci, suggesting – in agreement with ChIP-seq results – that it binds multiple genomic loci. As male germ cells differentiate into spermatocytes and prepare for meiosis, however, Rhi concentrates as one single dot in nuclei of early spermatocytes, suggesting its specific localization at one single locus. Since early male germ cells are diploid with two sets of autosomes, it is more plausible that this locus resides on one of the two sex chromosomes. To explore where Rhi binds at this stage, we first examined testes from XO males that lack Y chromosome and found that Rhi still localizes as one single dot in early spermatocytes, suggesting against Y-linked loci such as *Su(Ste)* and *h17* (Figure 6B). Simultaneous imaging of Rhi and RNA FISH of transcripts from the X-linked *AT-chX* locus revealed co-localization of *AT-chX* nascent transcripts and Rhi in one single dot from late spermatogonia to early spermatocytes, indicating that Rhi concentrates on *AT-chX* locus at this stage (Figure 6C). Indeed, single *AT-chX* RNA foci in individual nuclei became undetectable in *rhi* mutant testes, and expressing GFP-tagged Rhi transgene by *nanos-Gal4* in *rhi* mutant background restored *AT-chX* expression (Figure 6C). Importantly, even though *nanos-Gal4* drives stronger expression of GFP-Rhi in earlier stages (GSC and early spermatogonia), *AT-chX* transcripts remained highly expressed specifically during later stages (late spermatogonia and early spermatocyte) (Figure 6C). This finding suggests that the spatio-temporally regulated gene expression of *AT-chX* piRNA cluster is rather robust to perturbations to the Rhi protein level, and the low expression of *AT-chX* piRNA cluster earlier in GSCs and early spermatogonia is not limited by the level of Rhi protein. In sum, these results show that Rhi binds multiple genomic loci in GSCs and spermatogonia but concentrates on a single *AT-chX* locus later.

**Figure 6.**
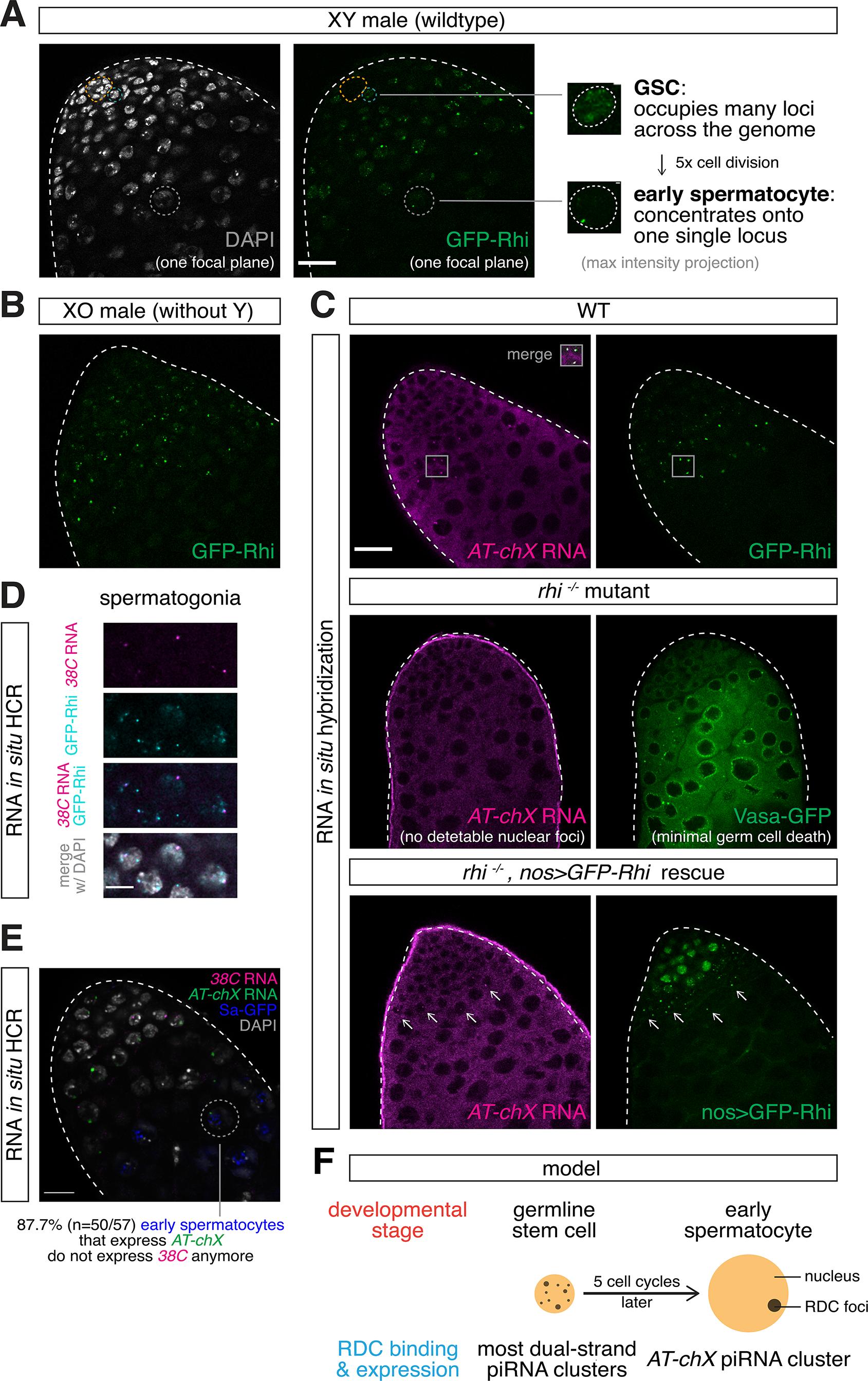
RDC complex enables dynamic piRNA production during early spermatogenesis. (A) Rhi localizes to multiple nuclear foci in GSCs and early spermatogonia but concentrates in a single dot in early spermatocytes. Confocal images of the apical tip of testis expressing GFP-Rhi under *rhi* promoter. A single focal plane is shown for DAPI and GFP-Rhi, while maximum intensity projections covering entire nuclei are shown for a GSC and an early spermatocyte on the right. Orange circle outlines the somatic hub for identification of GSCs next to it. Early spermatocyte is identified by formation of chromosome territory and increased nuclear size by DAPI. (B) Rhi localization to single nuclear dot is not affected in early spermatocytes that lack Y chromosome. Confocal image of the apical tip of testis from XO male (that lacks Y chromosome) expressing GFP-Rhi under *rhi* promoter. (C) Rhi localizes exclusively to *AT-chX* piRNA cluster in early spermatocytes. Top: RNA fluorescence *in situ* hybridization (FISH) of *AT-chX* piRNA precursors in wildtype testis expressing GFP-Rhi under *rhi* promoter. Middle: RNA FISH of *AT-chX* piRNA precursors in *rhi*^*2/KG*^ mutant testis. Note that the absence of *AT-chX* RNA foci is not due to germ cell death as Vasa expression shows minimal germ cell death. Bottom: expression of Rhi transgene driven by *nos-Gal4* rescues expression of *AT-chX* cluster in testis of *rhi*^*2/KG*^ mutant. (D) Rhi binds *38C* piRNA cluster in spermatogonia. RNA *in situ* hybridization chain reaction (HCR) of *38C* piRNA precursors in testis expressing GFP-Rhi under *rhi* promoter. Shown are spermatogonia as indicated by DAPI staining. Note that, in contrast to exclusive co-localization with *AT-chX* cluster in spermatocytes shown in (C), only a subset of Rhi foci co-localize with *38C* RNA foci, indicating that Rhi binds other piRNA clusters besides *38C* in spermatogonia. (E) *AT-chX* and *38C* co-express in spermatogonia, but only *AT-chX* is expressed in most early spermatocytes. Dual *in situ* HCR of *38C* and *AT-chX* piRNA precursors in testis expressing Sa-GFP (marker for spermatocytes). Circled is an example of early spermatocytes that express *AT-chX* but not *38C*. Quantification is shown at the bottom. Note that co-expression of *38C* and *AT-chX* can be seen in Sa-negative spermatogonia. (F) Proposed model of how Rhi switches genomic binding sites during gametogenesis from GSC to early spermatocyte stage, in order to allow dynamic employment of different piRNA clusters. All images share scale bars with (C), except for (D). Scale bars: 20µm (C) and 5µm (D).

As Rhi is required for non-canonical transcription of dual-strand piRNA clusters, depletion of Rhi from clusters other than *AT-chX* should cease their expression. To test this, we set out to conduct RNA FISH of piRNA precursors from other Rhi-dependent dual-strand piRNA clusters (Figure 5A). Previously, FISH detection of piRNA precursor transcripts was performed in polyploid nurse cells in fly ovary, whose genome is endo-replicated up to 1032C with a much higher expression level of piRNA precursor transcripts (Andersen et al., 2017; ElMaghraby et al., 2019; Mohn et al., 2014). In addition to low expression levels, piRNA cluster transcripts are difficult to target, as they are highly repetitive and share extensive sequence homology with TEs. To tackle these challenges, we employed *in situ* hybridization chain reaction (HCR), which permits enzyme-free signal amplification and automatic background suppression (Choi et al., 2018), to target transcripts from *38C* piRNA cluster that has a relatively high expression level in testis (Figure 5A). *in situ* HCR detected nascent *38C* piRNA precursors that co-localized with Rhi in diploid male germ cells (Figure 6D). Both *38C* and *AT-chX* piRNA precursors can be seen in nuclei of spermatogonia. In contrast, only *AT-chX*, but not *38C*, cluster continues to be expressed in 87.7% (n=50/57) of early spermatocyte nuclei (Figure 6E). Taken together, our results suggest that RDC complex binds, and thus enables, the expression of many dual-strand piRNA clusters from GSCs to spermatogonia, including *38C*, but gradually concentrates onto a single locus, *AT-chX*, in early spermatocytes. Dynamic association of RDC with piRNA clusters during early spermatogenesis executes a fluid piRNA program to allow different piRNA clusters to be engaged at different developmental stages.

## DISCUSSION

Most proteins involved in piRNA pathway in *Drosophila* were initially identified in screens for female sterility or TE de-repression in ovaries (Berg and Spradling, 1991; Czech et al., 2013; Handler et al., 2013; Karpen and Spradling, 1992; Muerdter et al., 2013; Schüpbach and Wieschaus, 1991). However, the first described case of piRNA repression, silencing of *Stellate* by *Su(Ste)* piRNAs during spermatogenesis, indicates that piRNA pathway operates in gonads of both sexes (Aravin et al., 2001; Vagin et al., 2006). Many proteins involved in piRNA pathway in ovaries are also required for male fertility and *Stellate* silencing in testes, supporting the conservation of piRNA pathway machinery between sexes (Aravin et al., 2001; Li et al., 2009; Nagao et al., 2010; Pane et al., 2007; Schmidt et al., 1999; Stapleton et al., 2001; Tomari et al., 2004; Zhang et al., 2011). Notably, a few proteins stood out as exceptions: Rhi, Del and Cuff that form a complex to enable transcription of dual-strand piRNA clusters in ovary (Klattenhoff et al., 2009; Mohn et al., 2014; Zhang et al., 2014), for which no fertility defects or *Stellate* de-repression were observed in mutant males (Klattenhoff et al., 2009; Schüpbach and Wieschaus, 1991; Volpe et al., 2001). This suggested that molecular mechanisms controlling piRNA cluster expression in testis might be different from ovary. Our results, however, demonstrate that RDC complex is assembled in testis (Figure 2) and required for TE silencing (Figure 3) in male germline, indicating that the molecular machinery regulating piRNA cluster expression is conserved between sexes.

### piRNA pathway in *Drosophila* testis: beyond *Stellate* silencing

Genomic loci that encode ovarian piRNAs were systematically identified across the genome in several studies (Brennecke et al., 2007; Mohn et al., 2014). In contrast, piRNA studies on testes were mostly focused on a single locus, *Su(Ste)*, that encodes piRNAs to silence *Stellate*, and no systematic search for piRNA clusters in testis has been performed to date. We *de novo* identified piRNA clusters in testis (see accompanying manuscript), laying the foundation for broader understanding of piRNA biogenesis and function in male gonads. We found that RDC is essential for expression of all major dual-strand piRNA clusters in testis, with a remarkable exception of *Su(Ste)* (Figure 4). This explains the previous observation that *rhi* is dispensable for *Stellate* silencing (Klattenhoff et al., 2009). In contrast, many other piRNA pathway factors such as Aub, Ago3 and Zuc are involved in both *Stellate* and TE repression (Aravin et al., 2001; Li et al., 2009; Pane et al., 2007). Mutations in these genes cause dramatic disruption of spermatogenesis, often leading to complete male sterility. In comparison, *rhi* mutant males demonstrate milder fertility defects, suggesting that *Stellate* de-repression is the major cause of spermatogenesis failure in other piRNA pathway mutants. Indeed, previous studies demonstrated that *Stellate* de-repression induced by deleting the *Su(Ste)* locus alone, without global perturbations of piRNA pathway, disrupts spermatogenesis and causes male sterility (Hardy et al., 1984). Thus, the finding that RDC is dispensable for *Stellate* repression provides a unique opportunity to understand impacts of silencing other piRNA targets in the male germline.

Our results indicate that piRNA-guided repression plays a crucial role in spermatogenesis beyond *Stellate* silencing, as *rhi* mutant males show rapid fertility decline, germline DNA damage and severe loss of germline content including GSCs (Figure 3). These phenotypes are likely caused by TE de-repression and the resultant genome instability, though we also identified complex satellites and a host protein-encoding gene, SUMO protease *CG12717*/*pita*, as targets of piRNA silencing in testes (see accompanying manuscript). In the future, it will be important to disentangle the contributions of de-repressing different piRNA targets to spermatogenesis defects observed in testes of piRNA pathway mutants.

Even though the fertility of *rhi* mutant males is substantially compromised, they produce a small number of functional spermatozoa, at least when they are young (Figure 1). In contrast, females lacking *rhi, del* or *cuff* are completely sterile (Schüpbach and Wieschaus, 1991; Volpe et al., 2001). This distinction between the two sexes might result from differences in the TE threat faced by male and female germ cells. Different TE families are activated upon disruption of piRNA pathway in ovary and testis and, generally, there is a stronger TE threat in ovary (see accompanying manuscript). Differential TE de-repression in two sexes might be responsible for stronger defects in oogenesis at the absence of RDC. Alternatively, it might reflect differences in DNA damage response in two sexes. An egg is energetically more expensive to make than a spermatozoon. In line with this, DNA damage responses (activated by e.g., TE transposition) often arrest oogenesis to avoid wasting resources on a defective egg (Chen et al., 2007; Klattenhoff et al., 2007). Since oogenesis is usually shut down to attempt repair, incomplete oogenesis results in female sterility. In contrast, quality control mechanisms of spermatogenesis frequently kill unqualified germ cells (Yacobi-Sharon et al., 2013), without pausing the developmental program of those surviving ones. Because spermatogenesis permits a large number of germ cells to develop in parallel, even though the unqualified ones are killed, a few surviving germ cells might be able to complete sperm development. Together, differential TE threats coupled with distinct response strategies to DNA damage might underlie sex-specific sterility when RDC is lost.

### *Su(Ste)*: an RDC-independent dual-strand piRNA cluster free of RDC binding

*Su(Ste)* locus on Y chromosome is the most prolific source of piRNAs in testes (see accompanying manuscript). piRNAs are generated from both genomic strands of *Su(Ste)* repeats, making it akin to other dual-strand piRNA clusters (Aravin et al., 2001, 2004). However, our results showed that RDC is dispensable for expression of *Su(Ste)* piRNAs, while it is required for piRNA production from all other dual-strand clusters in testis (Figure 4A) and ovary (Mohn et al., 2014; Zhang et al., 2014). What might explain the ability of *Su(Ste)* to generate piRNAs in an RDC-independent fashion? RDC ensures transcription of piRNA precursors by suppressing their premature termination (Chen et al., 2016) and promoting non-canonical transcription initiation (Andersen et al., 2017). Interestingly, the structure of *Su(Ste)* locus is different from other dual-strand piRNA clusters, as it is composed of many almost identical, relatively short units, all of which are flanked by two strong promoters driving expression of both genomic strands (Aravin et al., 2001). Thus, the presence of canonical promoters flanking individual short units of *Su(Ste)* repeats might enable their expression without engaging RDC complex.

Consistent with a role in promoting piRNA precursor transcription, Rhi is enriched on chromatin of dual-strand piRNA clusters. Such a correlation between direct Rhi binding and Rhi-dependent piRNA expression was previously reported in ovary (Mohn et al., 2014). The function of RDC explains why piRNA production depends on its presence on chromatin; however, the molecular mechanism responsible for specific recruitment of RDC to piRNA clusters remained poorly understood. It has been shown that piRNAs expressed during oogenesis are deposited into the oocyte and play an important role in jump-starting piRNA biogenesis in the progeny (Brennecke et al., 2008; de Vanssay et al., 2012). Maternally supplied piRNAs were shown to induce deposition of Rhi on cognate genomic locus in the progeny (Le Thomas et al., 2014). Thus, piRNAs expressed from the cluster and RDC binding to the cluster seem to form a feed-forward loop: RDC is required for piRNA production, and piRNAs in turn guide deposition of Rhi on cognate genomic loci. piRNA-dependent Rhi deposition might be mediated by the nuclear Piwi protein that directs the establishment of histone H3K9me3 mark (Le Thomas et al., 2013; Rozhkov et al., 2013; Sienski et al., 2012; Wang and Elgin, 2011), which provides a binding site for Rhi chromo-domain (Le Thomas et al., 2014; Mohn et al., 2014). Importantly, Piwi- and piRNA-dependent Rhi recruitment seems to occur in a narrow developmental window during early embryogenesis. This was demonstrated by the observation that depleting Piwi during early embryogenesis is sufficient to perturb Rhi localization on piRNA clusters, while depleting Piwi during larval or adult stages does not change Rhi localization (Akkouche et al., 2017). Our finding that Rhi is localized to all dual-strand piRNA clusters in testis except *Su(Ste)* (Figure 4E) is compatible with an idea that Rhi binding to genomic loci in the zygote is guided by maternal piRNAs. Indeed, in contrast to most other dual-strand clusters that are active in the germline of both sexes, Y-linked *Su(Ste)* locus generates piRNAs only in males. Therefore, unlike other piRNAs, *Su(Ste)* piRNAs are not deposited into the oocyte, resulting in the inability to recruit Rhi to *Su(Ste)* in the progeny. It is interesting to compare *Su(Ste)* with another dual-strand piRNA cluster on Y chromosome, *h17*. Unlike *Su(Ste)*, Rhi is enriched on *h17* chromatin and piRNA production from this cluster depends on Rhi (Figure 4A and 4E). However, *h17* piRNAs derived from male-specific Y should be absent in ovaries and hence no *h17* piRNAs can be deposited into the oocyte. At the first glance, these observations argue against a possibility that maternal piRNAs guide Rhi deposition on *h17* cluster. However, unlike *Su(Ste)*, *h17* is enriched in sequences of different TEs. As a result, TE-mapping piRNAs produced from other clusters in ovary might be able to target *h17*. Overall, while our results did not directly address the mechanism of Rhi recruitment to specific genomic loci, they show that studying Rhi occupancy on Y-linked piRNA clusters provides a novel angle to study this problem.

### Dynamic organization of piRNA pathway during spermatogenesis

Our results showed that components of RDC complex are expressed exclusively in male germline during early stages of spermatogenesis, from GSCs to early spermatocytes (Figure 1; Figure 2). Interestingly, expression of Piwi and Ago3, two of the three PIWI proteins in *Drosophila*, is also restricted to the same developmental stages (Cox et al., 2000; Nagao et al., 2010). Piwi is required for piRNA-guided transcriptional silencing in the nucleus (Le Thomas et al., 2013; Rozhkov et al., 2013; Sienski et al., 2012; Wang and Elgin, 2011), while Ago3 is involved in heterotypic ping-pong cycle in cytoplasm (Brennecke et al., 2007; Li et al., 2009; Malone et al., 2009), indicating that these processes operate in the same cells that have RDC-dependent transcription of piRNA clusters. In contrast to RDC, Piwi and Ago3, expression of the third PIWI protein, Aub, continues through spermatocyte stage until meiosis (Nagao et al., 2010). How can developing male germ cells be protected when the piRNA pathway is greatly simplified? It is possible that the silencing network initiated by piRNA pathway factors earlier can self-sustain later. piRNAs produced with the help of RDC complex and Ago3-dependent heterotypic ping-pong could persist and continue to function through spermatocyte differentiation, as long as they load onto Aub.

Interestingly, the cessation of RDC, Piwi and Ago3 expression during spermatogenesis coincides with the mitosis-to-differentiation transition. This transition is accompanied by one of the most dramatic changes in gene expression programs, with the general transcriptional machinery replaced by the testis-specific ones like tTAF and tMAC (Beall et al., 2007; Hiller et al., 2004). Also, after transition to spermatocytes, germ cells lose their ability to de-differentiate back to germline stem cells (Brawley and Matunis, 2004). It is thus tempting to propose that piRNA pathway contributes to the maintenance of cellular plasticity in early male germ cells (GSCs, gonialblasts and spermatogonia) by ensuring robust genome defense.

Restriction of the piRNA pathway to early stages of spermatogenesis contrasts with its activity during oogenesis. piRNA pathway factors appear to be expressed during all stages of oogenesis from GSCs to late-stage nurse cells. Recent studies reported several new factors involved in piRNA pathway in ovary, such as Moon, Boot, Nxf3, Panx, Arx and Nxf2 (Andersen et al., 2017; Batki et al., 2019; Dönertas et al., 2013; ElMaghraby et al., 2019; Fabry et al., 2019; Kneuss et al., 2019; Muerdter et al., 2013; Murano et al., 2019; Ohtani et al., 2013; Sienski et al., 2015; Yu et al., 2015; Zhao et al., 2019). These proteins are expressed at a low level in testis and their functions during spermatogenesis have not yet been reported. Our results suggest that, similar to RDC, these proteins might function in piRNA pathway in testis, and their low expression could be explained by restricted expression in early male germline. Indeed, we found that Moon co-expresses with, and acts genetically downstream of, RDC in testes (Figure 2; Figure S1). Overall, our results suggest that piRNA pathway machinery is likely conserved between sexes. However, the developmental organization of piRNA pathway during gametogenesis is different: whereas the entire pathway is active throughout oogenesis, processes that require RDC, Piwi or Ago3 likely terminate when mitotic spermatogonia differentiate into spermatocytes and prepare for meiosis.

### Dynamic expression of piRNA clusters during spermatogenesis

Our results revealed dynamic association of RDC complex with different genomic loci during spermatogenesis (Figure 6). In GSC and spermatogonia nuclei, RDC complex localizes to many foci, and ChIP-seq data indicate that Rhi associates with multiple dual-strand clusters. As germ cells differentiate into early spermatocytes, however, RDC gradually concentrates onto a single locus, *AT-chX*, on X chromosome. Since transcription of dual-strand piRNA clusters is dependent on RDC complex, the dynamic localization of RDC suggests that expression of piRNA clusters changes as germ cells progress from GSCs to early spermatocytes. Indeed, detection of nascent cluster transcripts revealed that *38C* is active early, but not later when most RDC concentrates onto *AT-chX.* Notably, though not dependent on Rhi, transcription of *Su(Ste)* piRNA cluster was shown to span a narrow window from late spermatogonia to early spermatocyte as well (Aravin et al., 2004), likely reflecting the promoter activity that drives *Su(Ste)* transcription (Aravin et al., 2001). Dynamic expression of piRNA clusters during spermatogenesis is also supported by the study that showed spermatogonia and spermatocytes have distinct piRNA populations (Quénerch’du et al., 2016). In contrast to dynamic localization of RDC complex and cluster expression in testes, previous work depicted a static view of piRNA production in female gonads. There has been no evidence of dynamic localization of RDC complex to different clusters during oogenesis, and all clusters appear active throughout female germline development. It will be interesting to explore whether expression of piRNA clusters changed dynamically during oogenesis. Through studying a protein complex thought to be absent during spermatogenesis, we uncovered the sexually dimorphic and dynamic behaviors of a molecular machinery that drives dual-strand piRNA cluster expression during *Drosophila* gametogenesis.

## ACKNOWLEDGEMENT

We are grateful to Xin Chen, Peter Andersen, William Theurkauf, Trudi Schüpbach, Paul Lasko, Elena Pasyukova and three Drosophila Stock Centers (Bloomington, Vienna, Kyoto) for fly stocks. We thank Katalin Fejes Toth and members of Aravin lab for discussion and comments. We appreciate the help of Maria Ninova and Fan Gao (Bioinformatics Resource Center, Caltech) with bioinformatics analysis, the help of Grace Shin and Maayan Schwarzkopf with HCR experiments, the help of Giada Spigolon and Andres Collazo (Biological Imaging Facility, Caltech) with microscopy, and the help of Igor Antoshechkin (Millard and Muriel Jacobs Genetics and Genomics Laboratory, Caltech) with sequencing. This work was supported by grants from the National Institutes of Health (R01 GM097363) and by the HHMI Faculty Scholar Award to A.A.A.

## SUPPLEMENTARY ITEM

**Figure S1.**
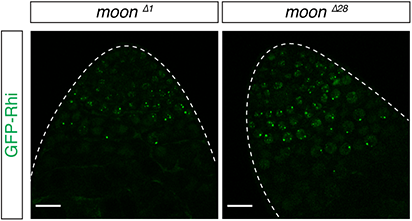
Moon acts downstream of Rhi in testis. Confocal images of apical tips of testes expressing GFP-Rhi, under *rhi* promoter, in *moon* ^∆*1*^ and *moon* ^∆*28*^ mutant backgrounds. Note that *moon* is X-linked, so XY males only have one copy of *moon* and trans-heterozygous mutant cannot be generated. Mutant alleles are nevertheless confirmed by causing female sterility (data not shown).

## MATERIALS AND METHODS

### Fly stocks

The following stocks were used: *bam*^∆*86*^ (BDSC5427), *bam*^*Df*^ (BDSC27403), C(1)RM (BDSC9460), *nos-Gal4* (BDSC4937), *UASp-shRhi* (BDSC35171), *iso-1* (BDSC2057) were obtained from Bloomington Drosophila Stock Center; *GFP-Rhi* (VDRC313340), *GFP-Del* (VDRC313271), *GFP-Cuff* (VDRC313269), *moon*^∆*1*^ (VDRC313735), *moon*^∆*28*^ (VDRC313738)*, Bam-GFP* (VDRC318001) were obtained from Vienna Drosophila Resource Center; *tj-Gal4* (DGRC104055) was obtained from Kyoto Stock Center; *rhi*^*2*^, *rhi*^*KG*^ and *UASp-GFP-Rhi* were gifts of William Theurkauf; *del*^*WK36*^, *del*^*HN56*^, *cuff*^*QQ37*^ and *cuff*^*WM25*^ were gifts of Trudi Schüpbach; *GFP-Vasa* (gift of Paul Lasko), *copiaLTR-lacZ* (gift of Elena Pasyukova), *GFP-Moon* (gift of Peter Andersen), *Sa-GFP* (gift of Xin Chen). *UASp-mKate2* was described before (Chen et al., 2016). XO males were generated by *GFP-Rhi* males crossed to C(1)RM females. *bam*^∆*86*^ and *GFP-Rhi* were recombined for testis ChIP.

### Generation of transgenic flies

To make mKate2-tagged Rhi driven by endogenous *rhi* promoter, the ~2Kb region upstream of *rhi* gene that includes the endogenous *rhi* promoter was cloned from genomic DNA of *Drosophila melanogaster* by PCR (forward primer: AGGCCTATGTACCAAGTTGTTAACTCTATCG, reverse primer: GGTACCAGACATAACTTATCCGCTCACAGG). PCR product was digested by Stu1 and Kpn1, then ligated into Stu1- and Kpn1-digested vector that contains mKate2-Rhino, mini-white gene and the ΦC31 attB site. The construct was inserted into genomic site 76A2 (y1 w1118; PBac{y+-attP-9A}VK00013) on chromosome 3 using ΦC31-mediated recombination.

### Sperm exhaustion test

The test was modified from Sun et al. (2004) and done at 25°C. Individual 1-day old virgin males (either *rhi*^*2/KG*^ or heterozygous siblings, n=15) were allowed to mate with three 4-day old wildtype virgin females (*iso-1*) for 24hrs. Each male was then moved to mate with another three 4-day old wildtype virgin females (*iso-1*) for another 24hrs, and this was repeated every 24hrs for a total of 14 days. Since each male encountered multiple young virgin females every day, their sperm were exhausted, and the number of progenies produced by females can be used to represent the daily male fertility. Inseminated females were flipped every other day and kept for 20 days (without contact with other males) to achieve maximal egg laying. A binary result of whether there is offspring or not was used to approximate whether a male is fertile or not on a given day, and we plotted the percentage of fertile male out of 15 tested males each day for *rhi*^*2/KG*^ and heterozygous siblings. To probe the male fertility more quantitatively, we repeated the test with five males for ten days. Instead of recording a binary result, we counted the number of progenies. The number of adult offspring was counted 15 days after female fly removal to allow most laid eggs to develop into adulthood. The averaged total number of offspring for each male each day was plotted for each genotype.

### Immunofluorescence staining

Testes were dissected in PBS, fixed in 4% formaldehyde for 20mins and washed by PBSTw (PBS with 0.1% Tween-20) for 3 times. Permeabilization of testes was done by incubation with PBST (PBS with 0.5% Triton-X) for 30mins. Testes were then blocked by 5% BSA in PBSTw for at least an hour, before incubation with primary antibody in 5% BSA in PBSTw at 4°C overnight. Testes were washed 3 times with PBSTw and incubated with secondary antibody in 5% BSA in PBSTw at room temperature for 2hrs, followed by another 3 washes with PBSTw. Before mounting in VECTA-SHIELD, testes were stained by DAPI (1:5000) for 10mins and rinsed once with PBS. The following primary antibodies were used: mouse anti-Fas3 (7G10, 1:200), mouse anti-γH2Av (UNC93-5.2.1, 1:400) and rat anti-Vasa (concentrated, 1:100) were obtained from Developmental Studies Hybridoma Bank.

### RNA fluorescence *in situ* hybridization (RNA FISH)

RNA FISH was done as described previously (Kotov et al., 2019). Fixed testes were prepared as above for immunofluorescence staining. Permeabilization was done by incubation in PBST with 0.5% sodium deoxycholate for an hour, followed by 3 washes of PBSTw. Testes were transferred to first 25% and then 50% formamide, both for 10mins. Next, testes were prehybridized in hybridization buffer (50% formamide, 0.5mg/ml yeast tRNA, 0.2mg/ml) for 1hr at 42°C, before incubation with 0.1-1µg DIG-labeled RNA probe in 50µl hybridization buffer overnight at 42°C with shaking. Testes were rinsed twice with 50% formamide for 20mins at 42°C and then transferred to wash in PBSTw for 4 times. Subsequent blocking, staining by sheep anti-DIG antibody (PA1-85378, 1:200, Life Technologies) and mounting were the same as described above for immunofluorescence staining. DIG-labeled RNA probes were transcribed by T7 according to manufacturer’s instructions. DNA template was made from genomic PCR using primers listed below, with T7 promoter sequence added 5’ to the reverse primers.

*Su(Ste)* (Aravin et al., 2004)

F: 5’-CAGGTGATTACCACTATTAACGAAAAGTATGC

R: 5’-ATCCTCGGCCAGCTAGTCCT

*AT-chX* (Kotov et al., 2019)

F: 5’-AGCGATCCCACTGCTAAAGA

R: 5’-ATAAAAGGTGACCG-GCAACG

### RNA *in situ* hybridization chain reaction (HCR)

A kit containing a DNA probe set, a DNA probe amplifier and hybridization, amplification and wash buffers were purchased from Molecular Instruments (molecularinstruments.org) for *AT-chX* and *38C* transcripts. To minimize off-targets, we designed probes targeting unique regions at *AT-chX* and *38C.* For *38C*, we specifically targeted junction sites of two different TEs, the simultaneous presence of which is required to generate amplified HCR signals. The *AT-chX* (unique identifier: 3893/E038) and *38C* (unique identifier: 4026/E138-E140) probe sets initiated B1 (Alexa647) and B3 (Alexa 546) amplifiers, respectively. *In situ* HCR v3.0 (Choi et al., 2018) was performed according to manufacturer’s recommendations for generic samples in solution.

### X-gal staining

Testes were dissected in PBS, fixed in 0.5% glutaraldehyde containing 1mM MgCl_2_ for 5mins and washed twice in PBS. Testes were incubated with 0.02% X-gal in X-gal buffer (1mM MgCl_2_, 150mM NaCl, 10mM Na_2_HPO_4_, 10mM NaH_2_PO_4_, 3.5mM K_4_Fe(CN)_6_ and 3.5mM K_3_Fe(CN)_6_) at 37°C in dark for time of interest. Staining of *copiaLTR-lacZ* in *rhi*^*2/KG*^ typically took 1.5-2.5hrs to develop. Reaction was then stopped by two washes of PBS and mounted as above for RNA FISH.

### Image acquisition and analysis

Images were acquired using confocal microscope Zeiss LSM 800 with 63x oil immersion objective (NA=1.4) and processed using the software Fiji (Schindelin et al., 2012). X-gal stained testes were imaged with 10x objective (NA=0.3). Maximum-intensity z-projection was done in Fiji, and line intensity profiles were obtained in Fiji. All images shown were from single focal planes, unless otherwise stated. Dotted outlines were drawn for illustration purposes. To quantify the number of GSCs, we stained the hub by Fas3. Z-stacks were acquired with 0.5µm intervals to cover depths well above and below the entire hub. Vasa-positive germ cells directly adjacent to the hub in 3D were deemed as GSCs and manually counted for each testis. Even though a molecular marker for GSCs was not used, any bias in GSC counting should be shared by both *rhi*^*2/KG*^ and heterozygous sibling controls.

### RNA-seq and analysis

RNA was extracted from dissected testes of 0-3 days old *rhi*^*2/KG*^ and heterozygous sibling controls using TRIzol (Invitrogen). About 1µg RNA for each sample was subject to polyA+ selection using NEBNext Poly(A) mRNA Magnetic Isolation Module (NEB E7490) and then strand-specific library prep using NEBNext Ultra Directional RNA Library Prep Kit for Illumina (NEB E7760) according to manufacturer’s instructions. For each genotype, two biological replicates were sequenced on Illumina HiSeq 2500 yielding 25-33 million 50bp single-end reads. Reads mapped to *D. mel* rRNA were discarded by bowtie 1.2.2 allowing 3 mismatches (<2% across all polyA-selected samples). For TE analysis, rRNA-depleted reads were mapped to TE consensus from RepBase17.08 using bowtie 1.2.2 with -v 3 -k 1. Mapped reads were normalized to the total number of reads that can be mapped to dm6 genome. Consistency between biological replicates was confirmed by >0.98 correlation coefficient. For simplicity, reads mapped to LTR and internal sequence were merged for each LTR TE given their well correlated behaviors. Note that polyA-selection was done for TE quantification in order to exclude piRNA precursor sequences that are not poly-adenylated but share sequence homology with TEs.

### piRNA-seq and analysis

RNA extraction was done as above for RNA-seq. 19-30nt small RNAs were purified by PAGE (15% polyacrylamide gel) from ~1µg total RNA. Purified small RNA was subject to library prep using NEBNext Multiplex Small RNA Sample Prep Set for Illumina (NEB E7330) according to manufacturer’s instructions. Adaptor-ligated, reverse-transcribed, PCR-amplified samples were purified again by PAGE (6% polyacrylamide gel). Two biological replicates per genotype were sequenced on Illumina HiSeq 2500 yielding 15-20 million 50bp single-end reads. Adaptors were trimmed with cutadapt 2.5 and size-selected for 23-29nt sequences for piRNA analysis. 23-29nt reads that mapped to rRNA were discarded by bowtie 1.2.2 tolerating 3 mismatches (<30% in control samples). For TE analysis, 23-29nt small RNA reads were mapped and normalized as done for polyA+ RNA described above, with correlation coefficients between replicates all >0.94. Complex satellite-mapping small RNAs were analyzed similarly (with ovary data downloaded from GSE126578). For piRNA cluster analysis, we used piRNA clusters defined in the accompanying manuscript. 1Kb genomic windows in individual piRNA clusters were generated with bedtools v2.28.0, and the ones including highly expressed miRNA, snRNA, snoRNA, hpRNA or 7SL SRP RNA were excluded. Coverage over individual piRNA clusters were computed using the pipeline tolerating local repeats described in the accompanying manuscript. A pseudo-count of 1 was added before calculating log2 fold-change of *rhi* mutant over control.

### ChIP-qPCR, ChIP-seq and analysis

ChIP protocol was modified based on Le Thomas et al. (2014). For each biological replicate, 200 pairs of 0-2 days old testes or 100 pairs of 4-5 days old ovaries (yeast-fed for 3 days) were fixed in 1% formaldehyde for 10mins, quenched by 25mM glycine for 5mins and washed 3 times with PBS. Fixed testes were snap-frozen in liquid nitrogen and stored in −80°C before ChIP. Frozen testes were first resuspended in PBS and then washed in Farnham Buffer (5mM HEPES pH8.0, 85mM KCl, 0.5% NP-40, protease inhibitor, 10mM NaF, 0.2mM Na_3_VO_4_) twice. Testes were then homogenized in RIPA Buffer (20mM Tris pH7.4, 150mM NaCl, 1% NP-40, 0.5% sodium deoxycholate, 0.1% SDS, protease inhibitor, 10mM NaF, 0.2mM Na3VO4) using a glass douncer and a tight pestle. Sonication was done in Bioruptor (Diagenode) on high power for 25 cycles (30s on and 30s off). Sonicated tissues were centrifugated to obtain the supernatant. The supernatant was pre-cleared with Dynabeads Protein G beads (Invitrogen) for 2hrs at 4°C. 5% of the pre-cleared sample was set aside as the Input, while the rest was incubated with 5µl anti-GFP antibody (A-11122, Invitrogen) overnight at 4°C. The immune-precipitated (IP) sample was incubated with Dynabeads Protein G beads for 5hrs at 4°C to allow beads binding. After that, beads were washed 5 times in LiCl Wash Buffer (10mM Tris pH7.4, 500mM LiCl, 1% NP-40, 1% sodium deoxycholate), while the Input sample was incubated with 1µl 10mg/ml RNase A at 37°C for 1hr. Both IP and Input samples were incubated with 100µg proteinase K in PK Buffer (200mM Tris pH7.4, 25mM EDTA, 300mM NaCl, 2% SDS) first at 55°C for 3hrs and then at 65°C overnight. DNA was purified by phenol-chloroform extraction and the concentration was measured by Qubit. Four biological replicates of testis (*bam*^∆*86/Df*^, *GFP-Rhi*) ChIP were done and used in qPCR, with two randomly selected replicates sequenced. Two biological replicates of ovary ChIP (*GFP-Rhi*) were done and sequenced. ChIP-qPCR was normalized first to Input and then to a negative control region (free of Rhi binding in ovary according to Mohn et al. 2014) to obtain Rhi enrichment (primers listed below). ChIP DNA was subject to library prep using NEBNext ChIP-Seq Library Prep Master Mix Set for Illumina (NEB E6240) according to manufacturer’s instructions. Two biological replicates per sex were sequenced on Illumina HiSeq 2500 yielding 13-22 million 50bp single-end reads. Reads were mapped to the genome as described in the accompanying manuscript permitting local repeats. Coverage over piRNA clusters were computed and the enrichment of IP over input was calculated. Two biological replicates were consistent with a correlation coefficient >0.96, so the average enrichment was plotted.

*42AB* (Klattenhoff et al., 2009)

F: 5’-GTG GAG TTT GGT GCA GAA GC

R: 5’-AGC CGT GCT TTA TGC TTT AC

*flam* (Klattenhoff et al., 2009)

F: 5’-TGA GGA ATG AAT CGC TTT GAA

R: 5’-TGG TGA AAT ACC AAA GTC TTG GGT CAA

*Su(Ste)* (Aravin et al., 2001)

F: 5’-CTTGGACCGAACACTTTGAACCAAGTATT

R: 5’-GGCATGATTCACGCCCGATACAT

*AT-chX* (Kotov et al., 2019)

F: 5’-AGCGATCCCACTGCTAAAGA

R: 5’-GTCGAAGACGTCCAGAGGAG

*negative control for Rhi binding* (this study)

F: 5’-AAGAGCAGAGGGGCCAAATC

R: 5’-TCCAAGTCGGCTTCCCTTTC

### Genome-wide correlation between piRNA production and Rhi binding

This analysis was adapted from Mohn et al. (2014) with modifications. piRNA production and Rhi enrichment were computed for individual 1Kb windows tiling the entire dm6 genome. The average of two well-correlative biological replicates was used for this analysis. Only 1Kb windows having both ≥4 RPM piRNAs in controls and ≥30 RPM reads in IP samples were plotted. Rhi-dependent loci (RD loci) were defined as 1Kb windows showing ≥4-fold drop in piRNA production in *rhi* mutant testes, and the rest were treated as Rhi-independent loci (RI loci). Local regression was implemented with LOESS technique in python.

### Data visualization and statistical analysis

Most data visualization and statistical analysis were done in Python 3 via JupyterLab using the following software packages: numpy (Oliphant, 2015), pandas (McKinney, 2010) and altair (VanderPlas et al., 2018). Germ cell death, GSC loss and ChIP-qPCR were plotted in GraphPad Prism. Mann–Whitney–Wilcoxon test was done to compute p values for GSC loss. Multiple t-tests corrected for multiple comparisons by the Holm-Sidak method were done for Rhi ChIP-qPCR, using the uni-strand cluster *flam* known to be free of Rhi binding as a negative control. The UCSC Genome Brower (Kent et al., 2002) and IGV (Robinson et al., 2011; Thorvaldsdóttir et al., 2013) were used to conduct explorative analysis of sequencing data.

### Data and code availability

Sequencing data will be uploaded to NCBI SRA.

